## Supplementary_Material for "Quantifying evolutionary constraints on the intronic branch point sequence in the bovine and human genome"

Animal Genomics, ETH Zürich, Eschikon 27, 8315 Lindau, Switzerland

### Supplementary Figure 1 – Variation in triplet codons

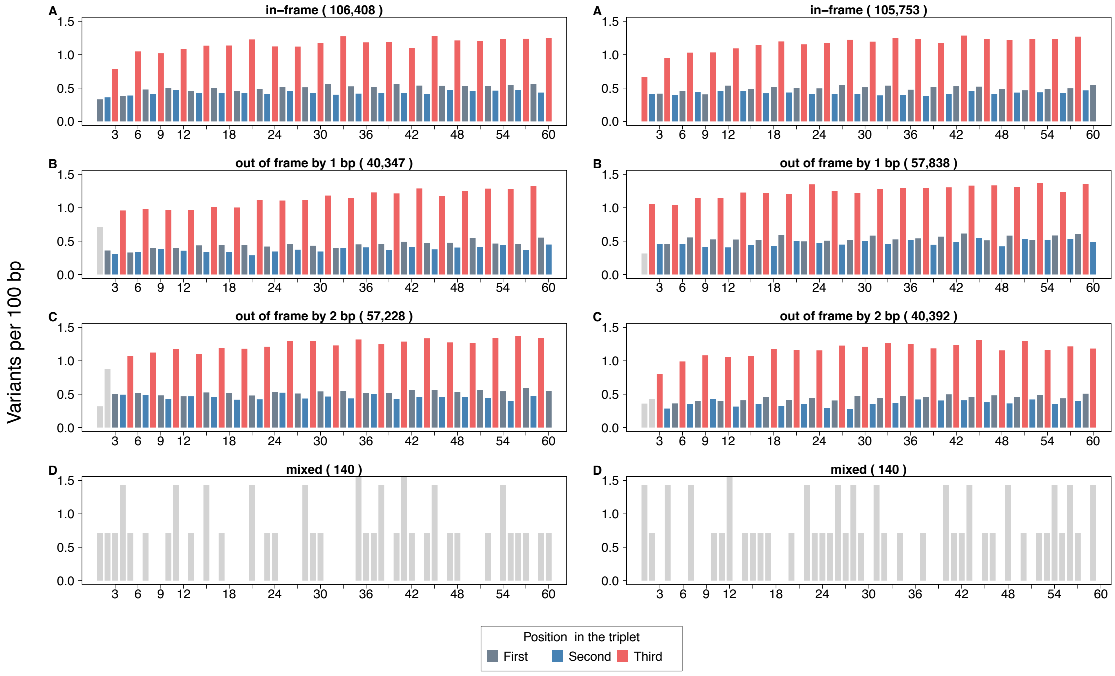

**Supplementary Figure 1**. Number of polymorphic sites (per 100 bp) overlapping the first, second and third base of coding triplets for exons starting with the first (A), second (B) and third (C) base of the triplet (left panel) and for exons ending with third (A), second. (B) and first (C) base of the triplet (right panel). The number of exons in each category is given in parenthesis in the plot title. Exons that do not consistently fall into the same category across transcripts of a gene are plotted in panel D. The frame information for exons were obtained from the gtf file.

#### Supplementary Figure 2 – Association between intron and BPS types

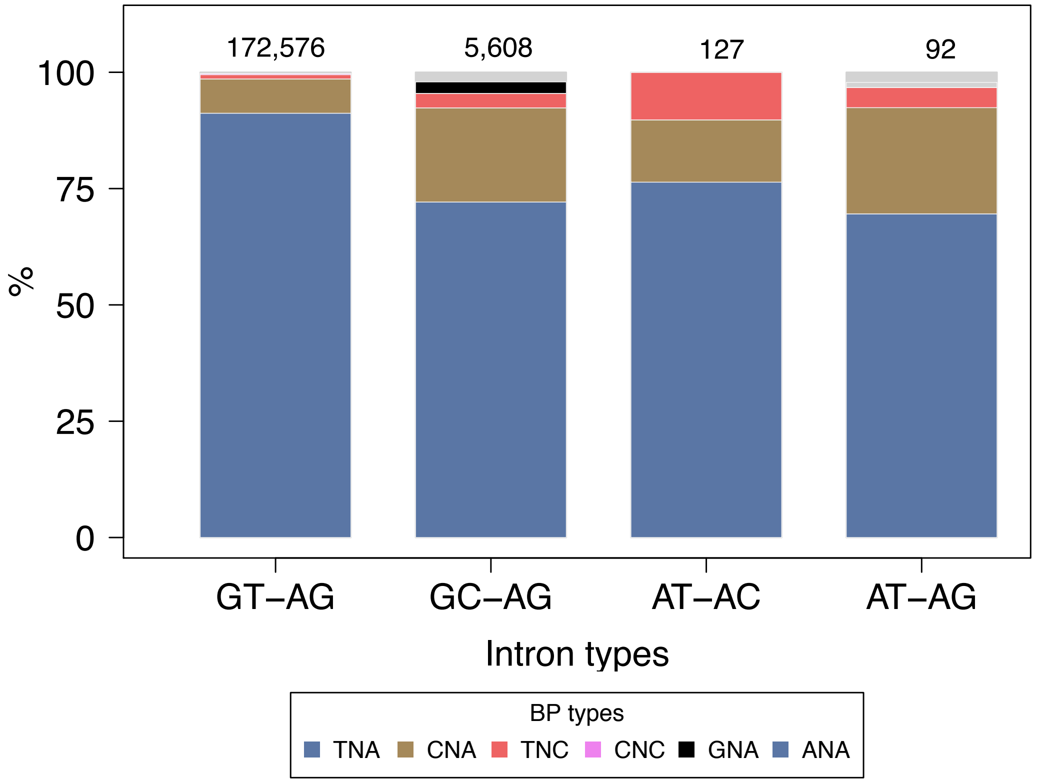

**Supplementary Figure 2.** Frequencies of intron types (5' ss – 3' ss) and the associated BPS (nucleotides at positions 4-6) in the bovine genome. The four most frequent intron types are represented as bars. Numbers on top of the bars indicate the occurrence of each intron type. The proportion of each branch point sequence type is indicated in different colors.

### Supplementary Figure 3 – Local variation in bovine branch point sequences predicted by LaBranchoR[1]

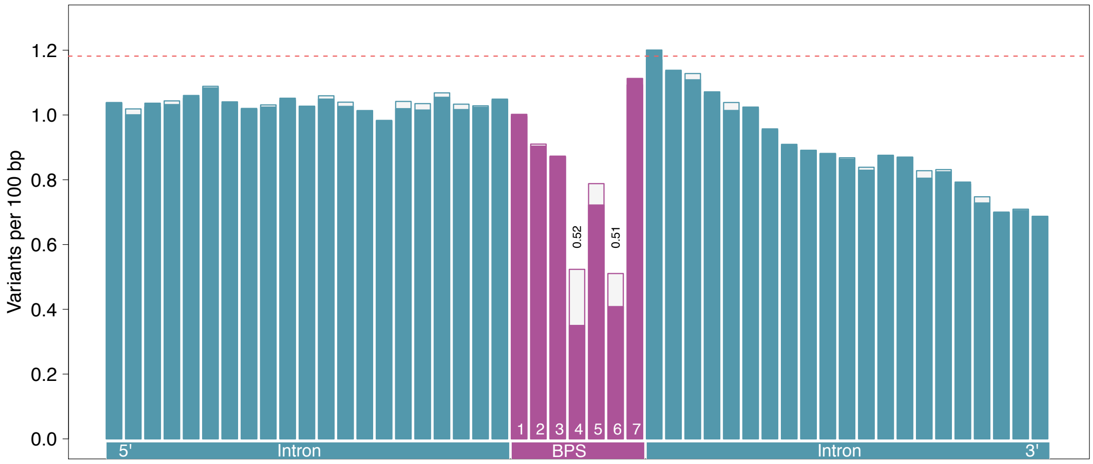

**Supplementary Figure 3**. Local variation at the bovine branch point sequence (BPS) predicted using LaBranchoR and 21 nucleotides on either side of the heptamer. The height of the bars indicates the variation (variants per 100 bp) surrounding 177,668 branch points while the solid bars indicate the variation surrounding a subset of 142,520 branch points encompassed by heptamers with a canonical (TNA) motif. The dotted horizontal red line indicates the average genome-wide variation.

#### Supplementary Figure 4 – Local variation in bovine branch point sequences predicted by branchpointer[2]

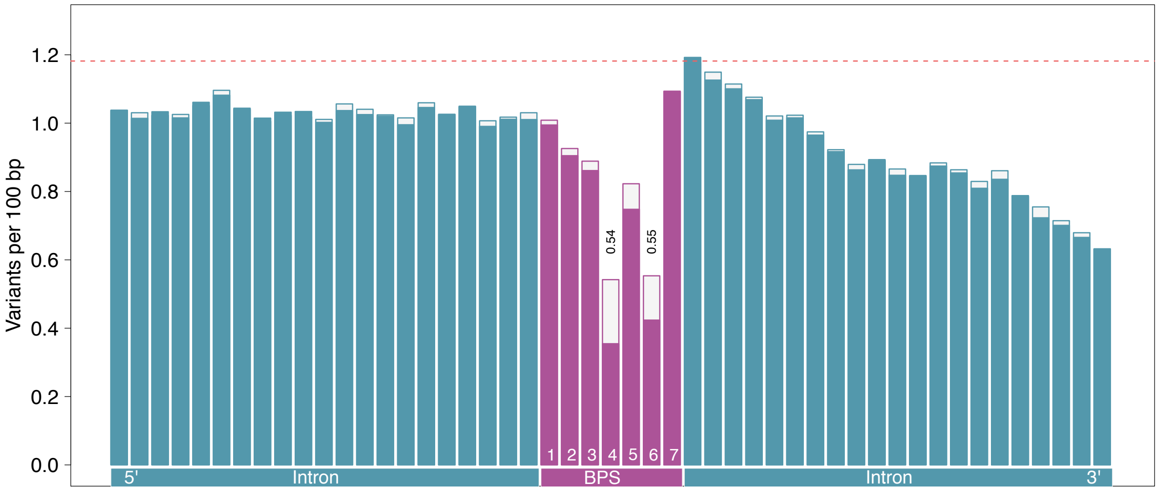

**Supplementary Figure 4**. Local variation at the bovine branch point sequence (BPS) predicted using branchpointer and 21 nucleotides on either side of the heptamer. The height of the bars indicates the variation (variants per 100 bp) surrounding 178,559 branch points while the solid bars indicate the variation surrounding a subset of 144,679 branch points encompassed by heptamers with a canonical (TNA) motif. The dotted horizontal red line indicates the average genome-wide variation.

### Supplementary Figure 5 – Local variation in a subset of bovine branch point sequences predicted at the same intronic position by BPP[3], LaBranchoR[1], and branchpointer[2]

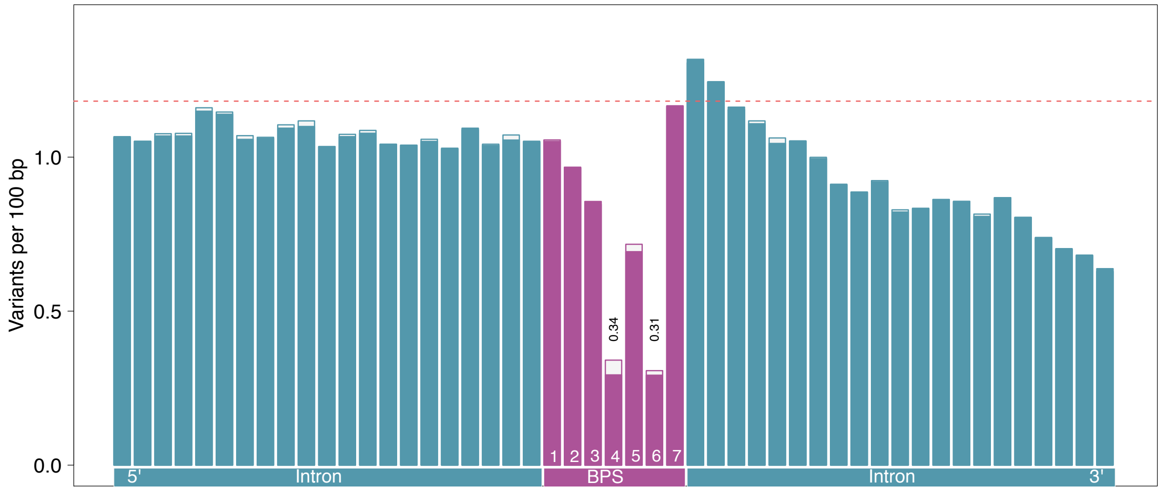

**Supplementary Figure 5.** Local variation around bovine branch point sequence (BPS) predicted using BPP, LaBranchoR, and branchpointer. The height of the bars indicates the variation (variants per 100 bp) surrounding 93,855 branch points predicted by all three tools while the solid bars indicate the variation surrounding a subset of 90,766 branch points encompassed within heptamers with a canonical (TNA) motif. The dotted horizontal red line indicates the average genome-wide variation.

### Supplementary Figure 6 – Local variation in a subset of human branch point sequences predicted at the same intronic position by BPP[3], LaBranchoR[1], and branchpointer[2]

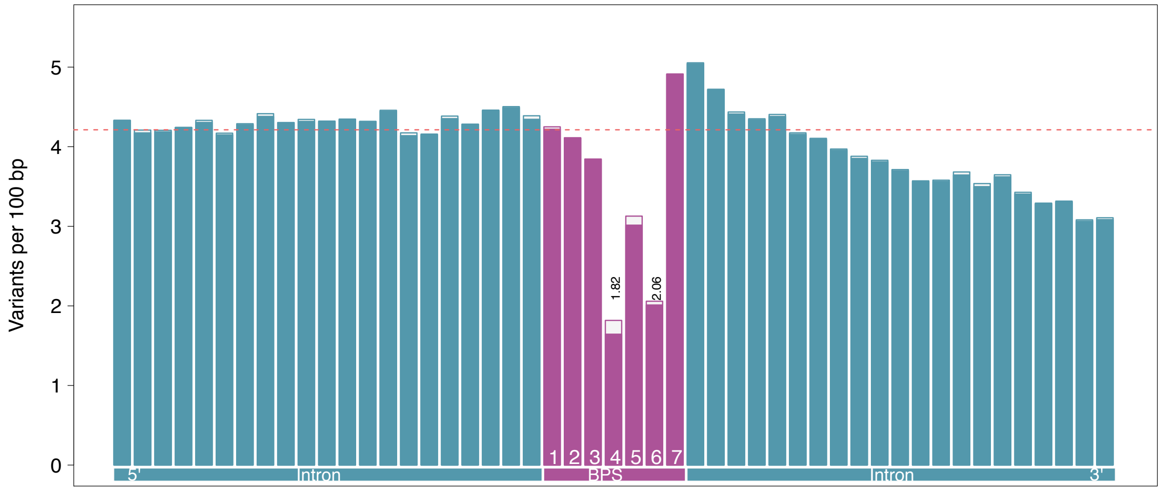

**Supplementary Figure 6.** Local variation around human branch point sequence (BPS) predicted using BPP, LaBranchoR, and branchpointer. The height of the bars indicates the variation (variants per 100 bp) surrounding 102,944 branch points predicted by all three tools while the solid bars indicate the variation surrounding a subset of 99,632 branch points encompassed within heptamers with a canonical (TNA) motif. The dotted horizontal red line indicates the average genome-wide variation.

### Supplementary Figure 7 – An exonic variant activates a cryptic splice site in bovine *WDR19*.

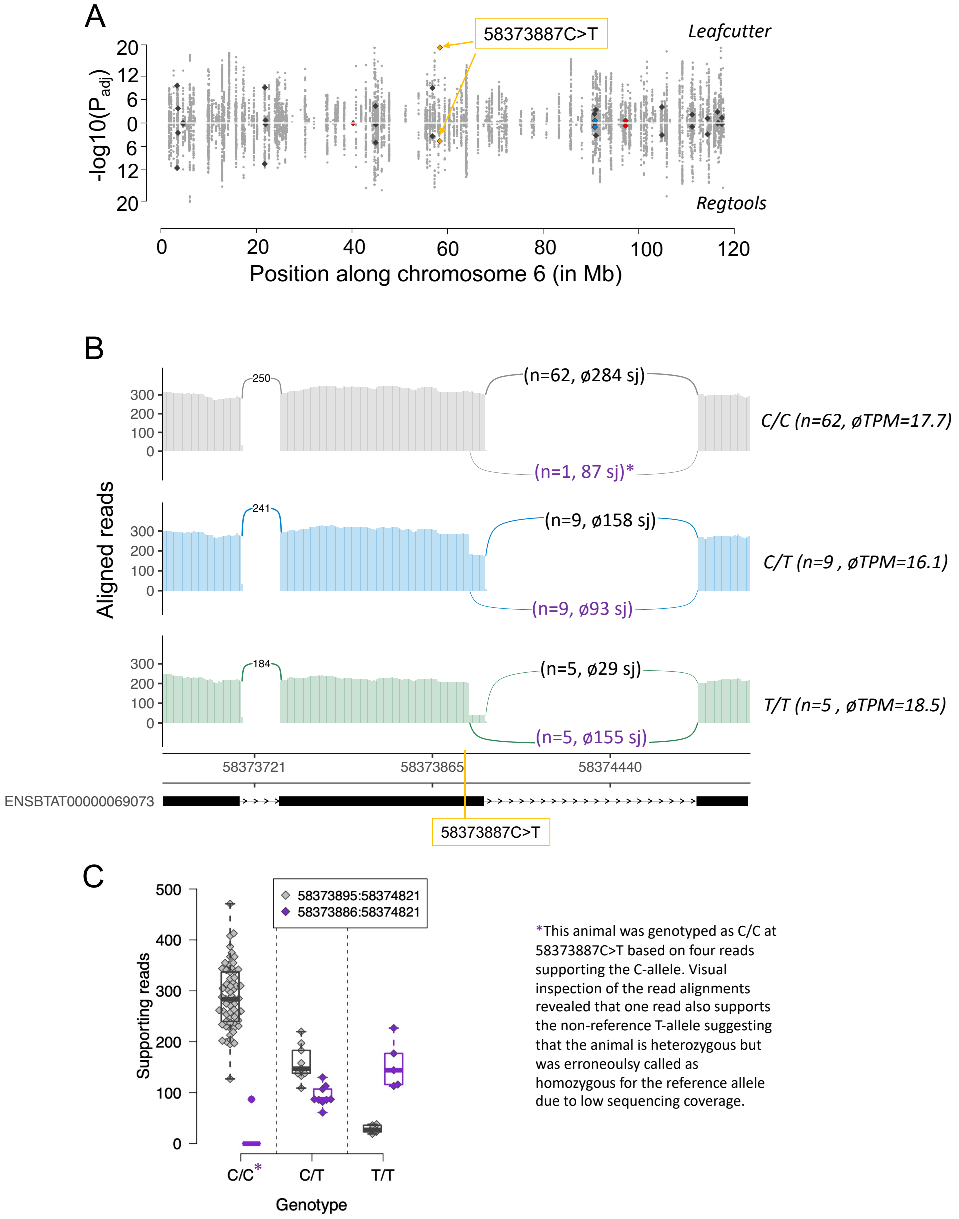

A) Manhattan plot of cis-acting splicing quantitative trait loci on bovine chromosome 6. Intron excision ratios were calculated based on exon junctions that were extracted from RNA sequencing read alignments using either Leafcutter or RegTools. Orange arrows indicate the BTA658373887C>T variant. This variant was reported to activate cryptic splicing, thus compromise male fertility. Red, blue, and dark-grey symbols represent variants affecting the fourth, sixth and all other positions of the heptamer, respectively. B) Sashimi plots of RNA sequencing coverage and splice junction (sj) utilization in 76 animals of the sQTL cohort with different BTA6:58373887 genotypes (grey: C/C, blue: C/T, green: T/T). Average abundance of *WDR19* mRNA in testis tissue is indicated in transcripts per million (TPM). Arcs indicate splice junction reads, with the thickness of the arc representing the average number of reads spanning the two exon junctions. Values in parentheses indicate the number of samples (n) for each genotype as well as the average number of junction-spanning reads. C) Boxplots and beeswarm plots of the number of reads supporting the primary (grey) and alternate (purple) splice junctions (sj).

### Supplementary Figure 8 – A mutation within a predicted branch point sequence is associated with alternative 3'splicing in FSD1L

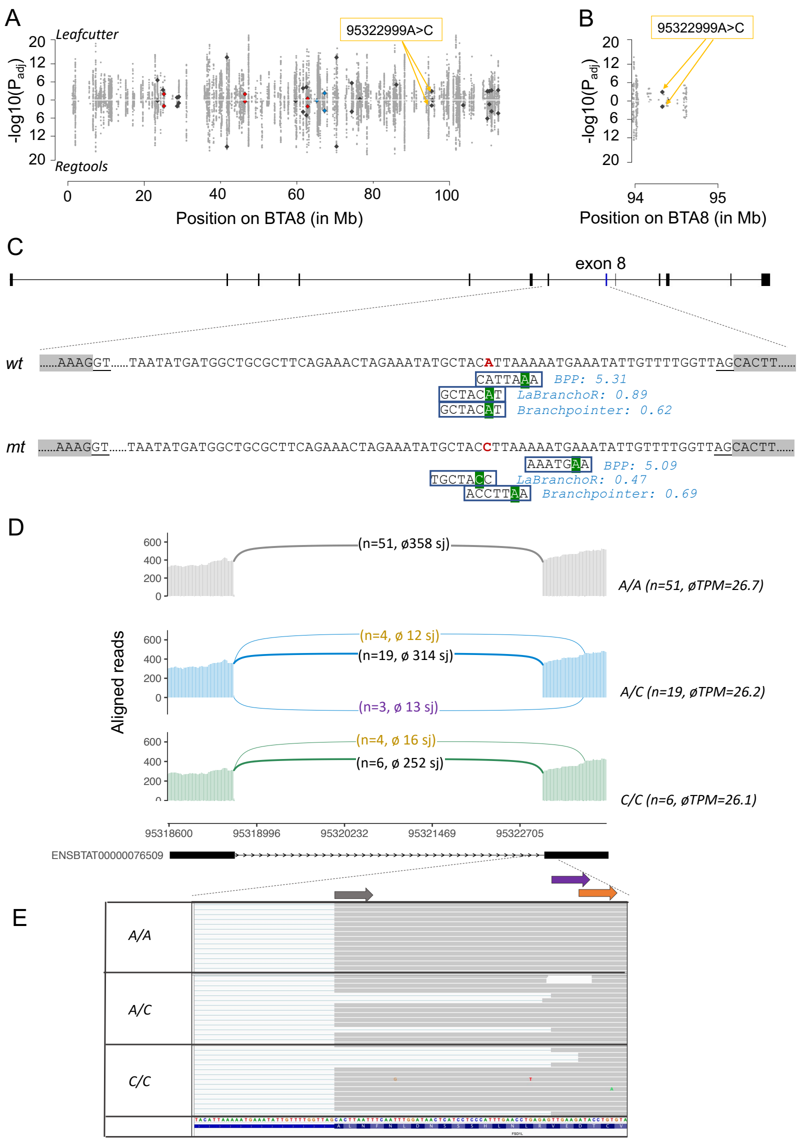

A) Manhattan plot of cis-acting splicing quantitative trait loci on chromosome 8. Intron excision ratios were calculated based on exon junctions that were extracted from RNA sequencing read alignments using either Leafcutter or RegTools. Orange arrows indicate the BTA8:95322999A>C variant. Red, blue, and dark-grey symbols represent variants affecting the fourth, sixth and all other positions of the heptamer, respectively. B) Detailed view of the region encompassing the BTA8:95322999A>C variant. C) Structure of the bovine *FSD1L* gene. Boxes represent exons. D) Branch point sequences predicted in the seventh intron of *FSD1L*. The BTA8:95322999 A-allele (red colour) constitutes the branch point predicted by LaBranchoR and branchpointer, and it is located at the seventh position of a heptamer encompassing a canonical «TNA»-motif that was predicted as the most likely branch point by BPP. Green background highlights the predicted branch point. Numbers reflect scores predicted by the different tools. The most likely branch point was placed at different locations for the intronic sequence with the BTA8:95322999 C allele. E) Sashimi plots of RNA sequencing coverage and splice junction (sj) utilization in 76 animals of the sQTL cohort with different BTA8:95322999 genotypes (grey: A/A, blue: A/C, green: C/C). Average abundance of *FSD1L* mRNA in testis tissue is indicated in transcripts per million (TPM). Arcs indicate splice junction reads, with the thickness of the arc representing the average number of reads spanning the two exons. Values in parentheses indicate the number of samples (n) for each genotype as well as the average number of junction-spanning reads. F)

Boxplots and beeswarm plots of the number of reads supporting the primary (grey) and two alternate (purple, orange) splice junctions (sj). G) Representative IGV screenshots of the RNA sequencing alignments for three animals with different BTA8:95322999A>C genotypes. Grey, purple, and orange arrows represent alternative exon starts.

### Supplementary Note 1 – Evolutionary constraints on functional features of the human genome

Using genotypes for 115,640,370 high quality SNPs segregating in 3,942 samples of diverse ancestry, we estimated the variability in different genomic features (Fig. 1). 488,127,421 bases overlapping 45,545 lncRNAs in the human genome were excluded from the intergenic region. On average, we detected 4.21 variants per 100 base pairs. Similar to findings in cattle, splice sites were the least variable positions in the genome.

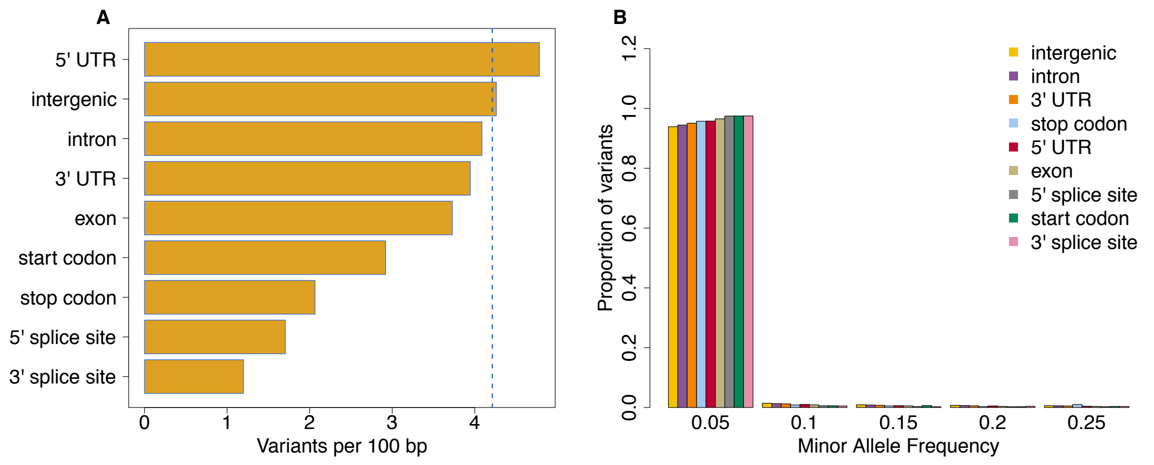

**Figure 1.** Variation (B) and allele frequency spectrum (C) within nine annotated genomic features. Variation is expressed as the number of variable nucleotides per 100 bp of the feature. The blue dotted line indicates the average genome-wide variation. Difference in variation was statistically significant between all pairs of features (Fisher's exact test $p<2.1 \times 10^{-13}$).

The overall variability of the features was consistent with findings in cattle except for the 5'UTR region, which was exceptionally variable (4.8; 14% more than the genome average) in humans. We observed a large standard deviation in the variability estimates of individual 5' UTRs, with smaller UTRs being more variable (Fig. 2). The local variation at the exon-intron boundaries also closely followed the pattern observed in the cattle data (Fig. 3).

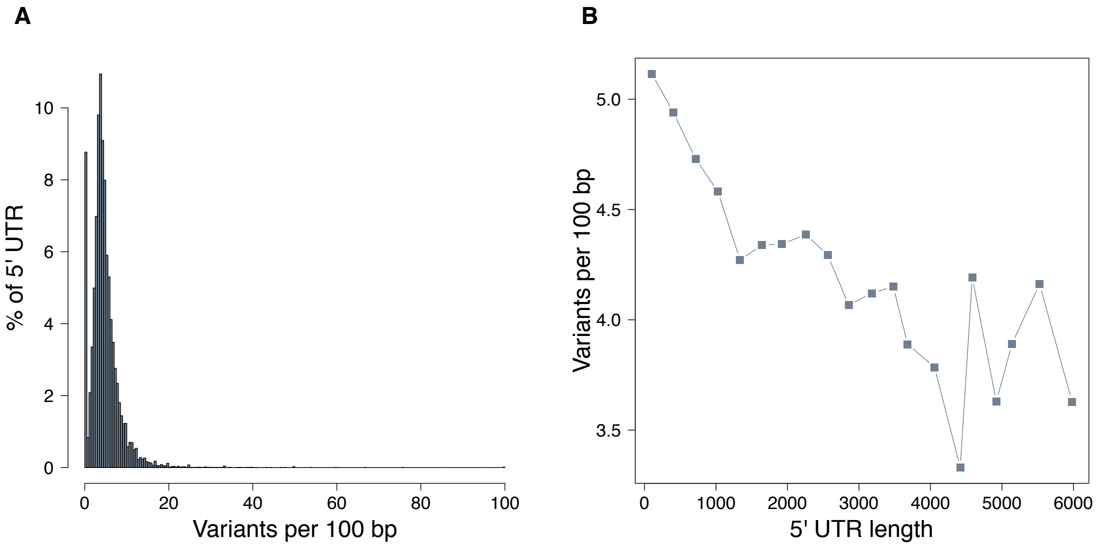

**Figure 2.** Distribution of variation (variants per 100 bp) in individual UTRs (A) and the relationship between UTR length and variation (variants per 100 bp) (B). Number of variants overlapping 100,948 5' UTR in the human genome was calculated. The UTRs were grouped into bins of 300 bases and mean variation was calculated for each bin.

We observed a general decline in variation at regions encompassing the splicing acceptor and donor bases with the strongest depletion of variation (68 and 54% less variation than exons) at the four nucleotides overlapping the splice sites. The periodic pattern of reduced variation in the third base of the codon triplet was also evident in the human data. These findings corroborate that the evolutionary constraint is stronger on splice sites than coding sequences.

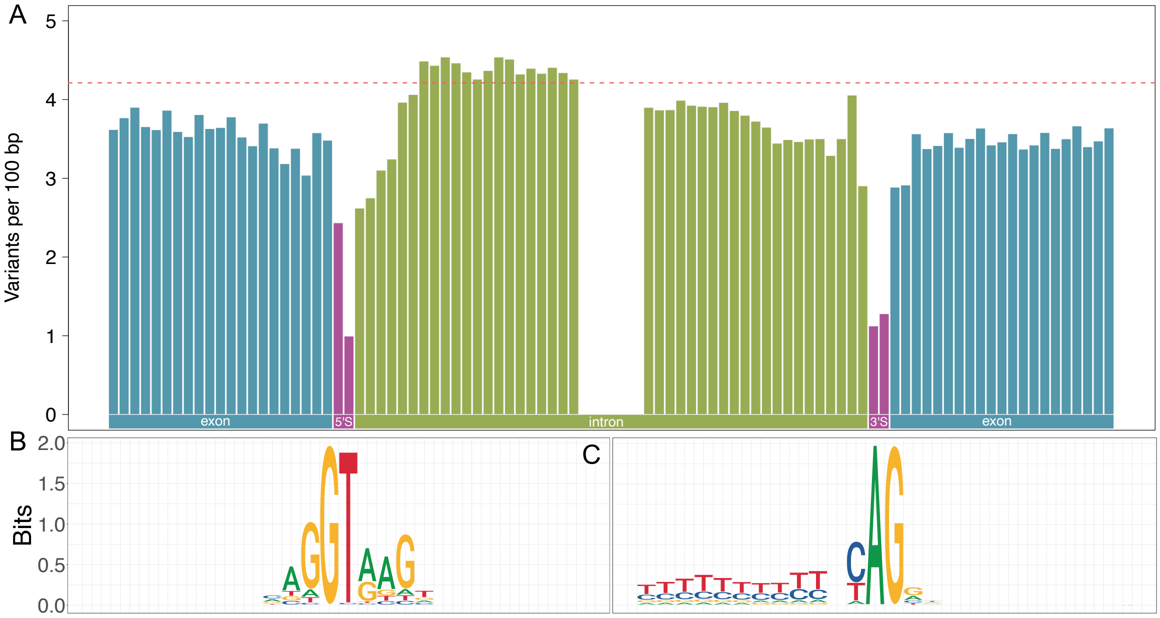

**Figure 3**. Evolutionary constraint around splice sites. Variation observed at 21 bases up- and downstream of the 3' and 5' splice sites (A). The red dotted line represents the average genome-wide variation. The conservation of nucleotides at 5' (B) and 3' (C) splice sites is shown in sequence logo plots.

#### Branch point sequence

The consensus branch point sequence was nnyTr<u>A</u>y (Fig. 4A; branch point underlined) and was placed between 14 and 145 (mean=29.3±11.9) bp upstream of 3' splice site (Fig. 4B). Like in cattle, the branch point was predominantly adenine (n=199,558, 98.87%) and position 4 of the heptamer was highly conserved with 92.33% of the predicted BPS carrying a thymine nucleotide at that position (Fig. 4A). A reduced Ti/Tv ratio at the branch point and position 4 of the heptamer was also observed in the human data (Fig. 5)

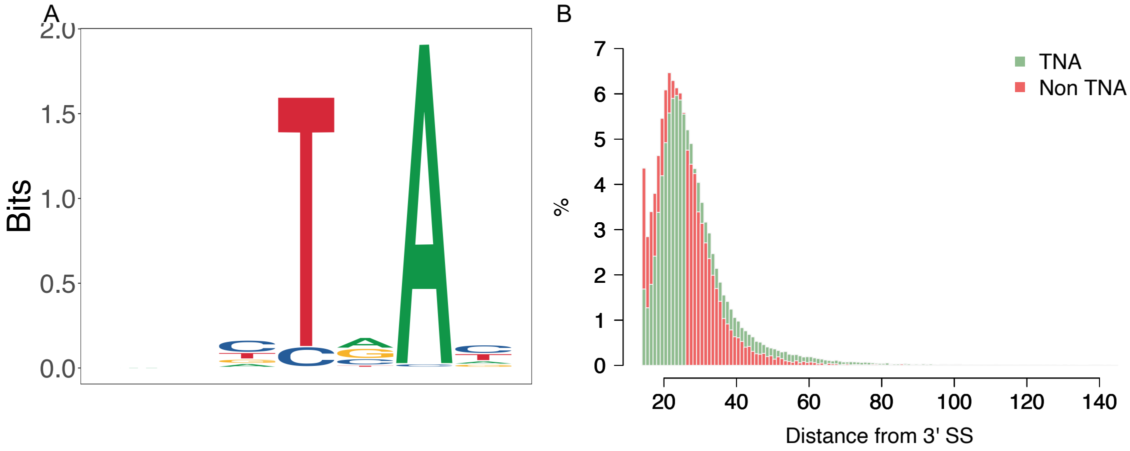

**Figure 4**: sequence logo plot generated from 201,832 branch point sequences identified in the human genome (A). Distribution of distance from 3' splice site to the predicted branch point adenine for canonical (green) and non-canonical (red) BPS (B).

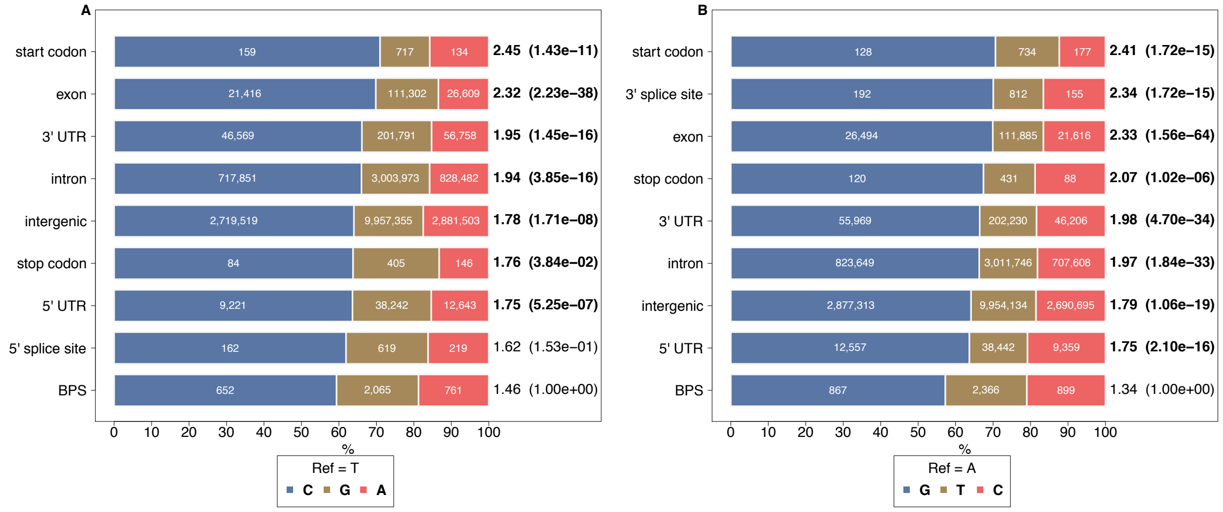

**Figure 5**. SNP mutation types of thymine at position 4 (a) and adenosine at position 6 (b) within canonical ("TNA") branch point sequence motifs and other genomic features.

The widths of differently colored bars indicate the proportion of substitution to the other three nucleotides. The number of observed substitutions is given in the middle of each differently colored bar. The Ti/Tv for the features is indicated at the end of the bars. The p values (<0.05 indicated in bold text) for the pairwise Fisher's exact test for Ti/Tv at the nucleotide in BPS vs nucleotides in other genomic features are given in parenthesis. For splice sites, analysis was limited to a splice site where the canonical sequence motif (GT and AG at 5' and 3' splice sites respectively) contains the studied residues.

#### Supplementary Note 2 – Comparison of branch point sequence prediction tools

We compared the branch point sequence (BPS) prediction from the program BPP to the predictions from the two other programs, LaBranchoR[1] , and branchpointer[2]. While BPP considers the entire intronic sequence of any length (we considered up to 250 bp upstream of the 3' splice site for introns longer than 20bp) to search for branch point sequences, the programs LaBranchoR and branchpointer restrict the branch point sequence search space to intronic sequences that are at least 70 bp long and 27 bases between 18 and 44 bases upstream of the 3' splice site, respectively.

Among the 179,476 unique intronic sequences considered for branch point sequence prediction using BPP, 177,668 and 178,559 introns also had predictions from LaBranchoR and branchpointer respectively. The predictions from LaBranchoR and branchpointer had the largest overlap of 75% (133,972/177,667) and the overlap in predictions was similar for BPP and LaBranchoR (60.3%; 108,251/177,668) and BPP and branchpointer (60.9%; 107,696/178559). For a subset of 177,667 introns that had predictions from all three tools, the predicted branch point was placed at the same locus for 93,855 (52.8%) introns (Fig. 1).

Next, we studied the distribution of scores for predicted BPS from the three tools. The mean score for 93,855 overlapping BPS was significantly higher than the overall mean score. The difference was only 0.4% for BPP (one tailed t-test; $p=2.7x10^{-06}$), however was >10% for the two other tools (one tailed t-test; $p < 1.0x10^{-300}$).

The scores from the tools were weakly but significantly negatively correlated with the variability of the predicted branch point residue (position 6 of the predicted heptamer). Heptamers predicted with higher confidence showed lower variability at the branch point. This correlation was the strongest for branchpointer and weakest for BPP (Fig. 2).

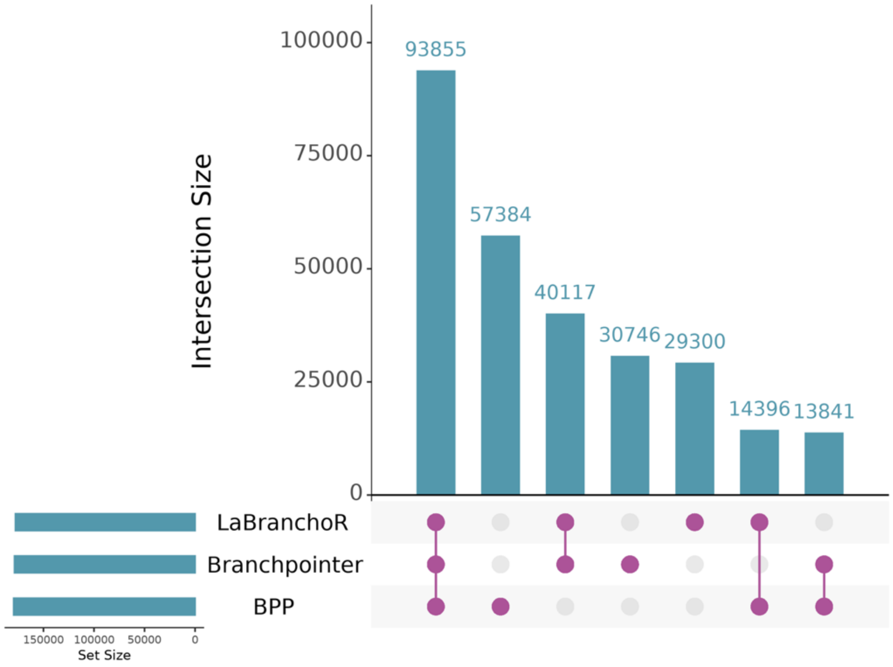

**Fig. 1**: UpsetR plot of 279,639 predicted unique branch point sequences in 179,476 introns from three prediction tools.

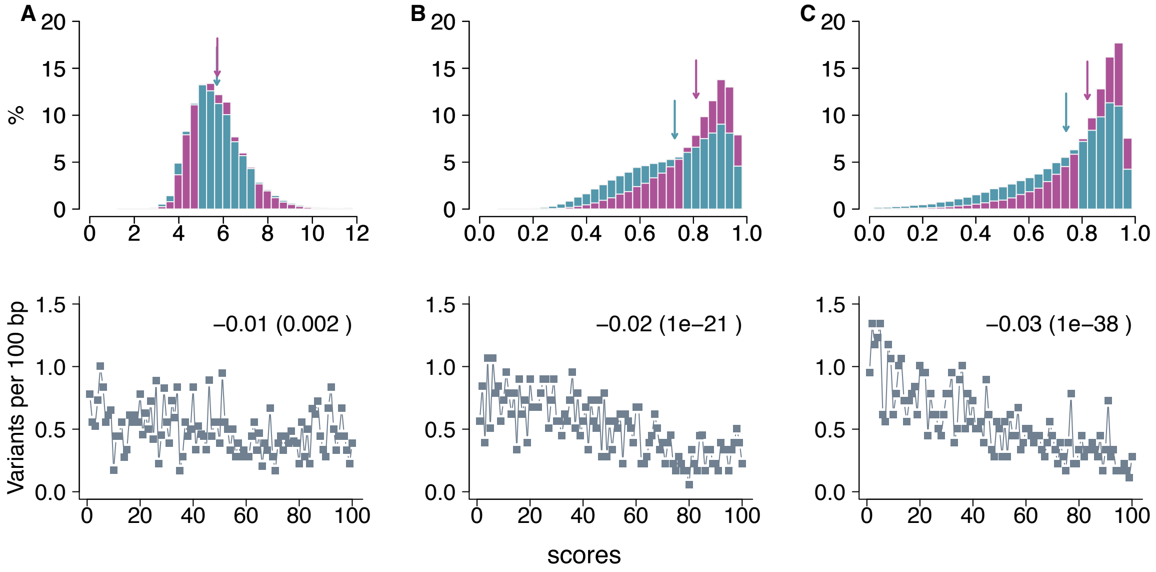

**Fig. 2:** Prediction scores (top panel) for all (blue) and 93,855 overlapping (pink) bovine branchpoints predicted using BPP (A), LaBranchoR (B) and, branchpointer (C). The colored arrows indicate the corresponding mean scores. Relationship between prediction scores (grouped into 100 bins) and variability of branchpoints (bottom panel) predicted using BPP (A), LaBranchoR (B) and, branchpointer (C). The correlation between prediction scores and branch point variability (variants per 100 bp) is given on the top right-hand corner with its statistical significance in the parenthesis.

Similar observations were made for branch point sequence prediction in human introns. Predictions were available, respectively for 201,832, 201,502 and 201,754 introns from BPP, LaBranchoR and branchpointer. For the 201,502 introns with predictions from all three tools, predictions overlapped for 102,944 (51.09%) of introns (Fig. 3). The overlap of predictions from BPP and the two other tools was ~60% as seen in the cattle data. The highest overlap, like in the cattle data, was observed for predictions from LaBranchoR and branchpointer (75%; 151,149/ 201,502).

The prediction scores from LaBranchoR and branchpointer were ~10% higher (two tailed t-test; $p<1.0x10^{-300}$) for the 102,944 introns with matching predictions from all three tools (Fig. 4). The scores from BPP did not differ significantly between all and matching predictions. The scores from LaBranchoR and branchpointer were also weakly but significantly negatively correlated with the variability of the predicted branchpoint, mirroring findings from the cattle data.

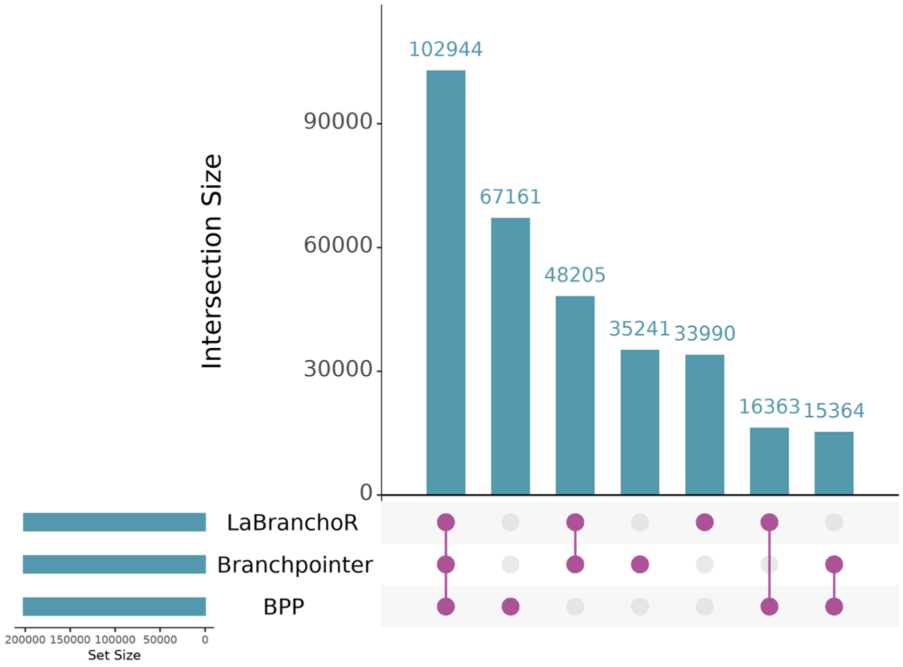

**Fig. 3**: UpsetR plot of 319,268 predicted unique human branch point sequences in 201,832 introns from three prediction tools.

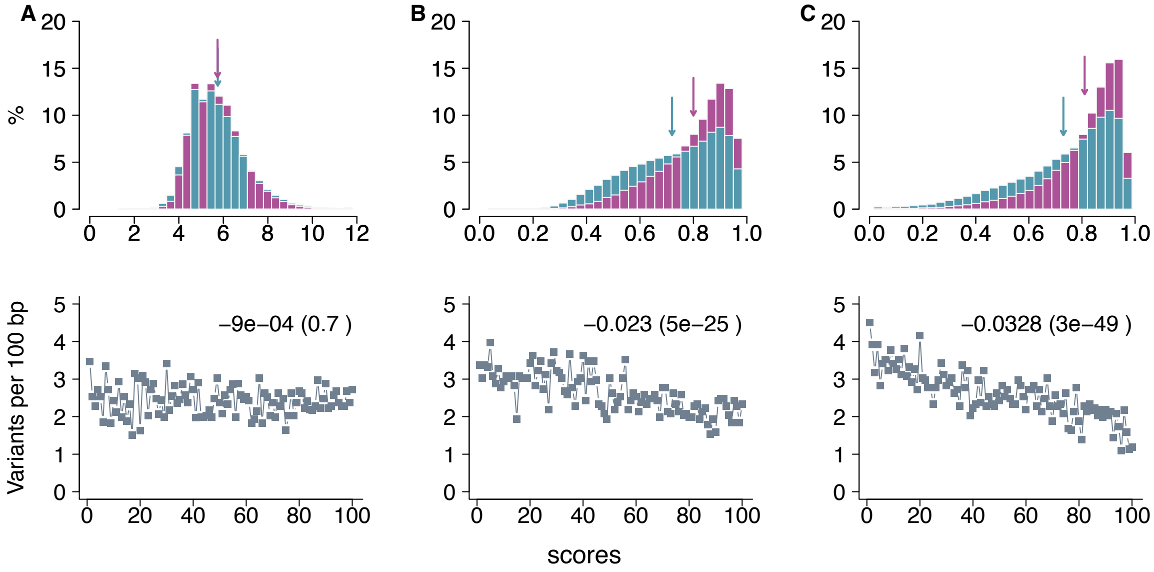

**Fig. 4:** Prediction scores (top panel) for all (blue) and 102,944 overlapping (pink) human branchpoints predicted using BPP (A), LaBranchoR (B) and, branchpointer (C). The colored arrows indicate the corresponding mean scores. Relationship between prediction scores (grouped into 100 bins) and variability of branchpoints (bottom panel) predicted using BPP (A), LaBranchoR (B) and, branchpointer (C). The correlation between prediction scores and branch point variability (variants per 100 bp) is given on the top right-hand corner with its statistical significance in the parenthesis.

#### Supplementary Note 3 – Mutations introducing novel AG dinucleotides in the AG exclusion zone of the bovine genome.

For all introns with canonical "AG" splicing site at the 3' splice site (n=178,690), we defined the AG exclusion zone (AGEZ) as the region between the end of the branch point sequence (predicted using BPP) and intron end excluding the splice sites. We then considered genomic positions of 185,703,494 adenine residues (778,931 in and 184,924,563 out of the AGEZ) and 135,839,698 guanine residues (632,125 in and 135,207,573 out of the AGEZ) that are not a part of AG dinucleotides in the reference sequence. All mutations next to these positions (one bp downstream of adenosine and one bp upstream of guanine) were tabulated using our variant catalogue of 29,227,937 biallelic variants. The rate of mutation was expressed per 100 bases considered and the statistical significance of difference in this rate in and out of the AGEZ was calculated using Fisher's exact test (Table 1).

The mutation rate was generally higher (~13%; in positions adjacent to adenine and guanine) outside the AGEZ (Fisher's exact test $p=8.1x10^{-59}$) corroborating our previously observed constraints nearby the 3' splice site (cf. Figure 2 in the main manuscript). However, mutations resulting in novel AG (i.e., mutation to guanine downstream of adenine and to adenine upstream of guanine) dinucleotides occurred more than twofold more often outside than inside the AGEZ. This difference was statistically highly significant for mutations adjacent both adenine and guanine residues (Fisher's exact test $p< 7.0x10^{-155}$).

**Table 1**. Number of mutations to the four nucleotides (Mut_adj_pos) adjacent to intronic adenine and guanine in (Num_in) and out (Num_out) of the AGEZ in the bovine genome. The rates of mutation in (Rate_in) and out (Rate_out) of the AGEZ are given along with the ratio (Rate_out/Rate_in) and significance from the Fisher's exact test. The mutations resulting in insertion of novel "AG" dinucleotides are given in bold.

| Base | Mut_adj_pos* | Num_in | Rate_in | Num_out | Rate_out | ratio | P |
|---:|---:|---:|---:|---:|---:|---:|---:|
| A | A | 818 | 0.11 | 303,529 | 0.16 | 1.45 | $4.7e^{-43}$ |
| A | C | 2,187 | 0.28 | 542,610 | 0.29 | 1.04 | $4.0e^{-02}$ |
| **A** | **G** | **1,108** | **0.14** | **532,546** | **0.29** | **2.07** | $\mathbf{1.8e^{-155}}$ |
| A | T | 2,837 | 0.36 | 612,678 | 0.33 | 0.92 | $6.7e^{-07}$ |
| **G** | **A** | **1,219** | **0.19** | **534,072** | **0.39** | **2.05** | $\mathbf{2.8e^{-177}}$ |
| G | C | 2,567 | 0.41 | 522,810 | 0.39 | 0.95 | $1.4e^{-02}$ |
| G | G | 781 | 0.12 | 255,440 | 0.19 | 1.58 | $3.7e^{-37}$ |
| G | T | 4,300 | 0.68 | 772,726 | 0.57 | 0.84 | $1.1e^{-25}$ |

*One bp downstream when the "Base" is A and one bp upstream when the "Base" is G

### References

1. Paggi, J. M. & Bejerano, G. A sequence-based, deep learning model accurately predicts RNA splicing branchpoints. *RNA* **24**, 1647–1658 (2018).
2. Signal, B., Gloss, B. S., Dinger, M. E. & Mercer, T. R. Machine learning annotation of human branchpoints. *Bioinformatics* **34**, 920–927 (2018).
3. Zhang, Q. *et al.* BPP: A sequence-based algorithm for branch point prediction. *Bioinformatics* **33**, 3166–3172 (2017).
